# Donor transcription suppresses D-loops in *cis* and promotes genome stability

**DOI:** 10.1101/2025.02.12.637806

**Authors:** Yasmina Djeghmoum, Aurèle Piazza

## Abstract

D-loops are DNA joint molecule intermediates central to DNA break repair by homologous recombination (HR). Priority rules between recombination and transcription at the donor locus have not been investigated. Here, using a controlled break induction system and physical detection of D-loops in *S. cerevisiae*, we show that donor transcription by RNA polymerase II acutely suppresses D-loops in *cis*, in an orientation-dependent manner. This inhibition does not rely on endogenous transcription factors, the RNA product, RNA:DNA hybrids, or previously characterized D-loop disruption factors. Transcription can be the major D-loop suppression pathway and inhibits the formation of repeat-mediated genome rearrangements. Transcription is therefore a negative regulator of HR at the D-loop level that promotes genome stability. These findings reveal the functional prioritization between two universal DNA-dependent processes.

**Highlights:**

- Donor transcription suppresses D-loops acutely and in *cis*
- Donor transcription is independent from conserved *trans* D-loop disruption pathways
- Transcription directionality modulates D-loop suppression
- Transcription locally enforces HR fidelity

## Introduction

Various protein-dependent processes such as transcription, replication, and recombination compete for the same DNA substrate and must be coordinated. Transcription has long been known to stimulate spontaneous HR in prokaryotes and eukaryotes by interfering with DNA replication, despite elaborate mechanisms that coordinate their deployment and mitigate conflicts (reviewed in (Aguilera & Gaillard, 2014; Goehring *et al*, 2023)). Comparatively much less is known about how transcription and homologous recombination (HR) are coordinated, and how putative priority rules impact genome stability.

HR is a high-fidelity DNA double-strand break (DSB) repair pathway that uses an intact homologous dsDNA molecule as template (Kowalczykowski, 2015). It entails the formation of a metastable DNA joint molecule called a Displacement loop (D-loop), which consists of a heteroduplex DNA (hDNA) region, a displaced strand and two non-identical 5’ and 3’ strand exchange junctions (Savocco & Piazza, 2021). Their extension by a DNA polymerase commits to repair while their disruption reinitializes the search for a homologous donor and eliminates toxic joint molecules. This reversibility, enforced by several conserved ancillary HR factors (Sgs1-Top3-Rmi1^BLM-TOPO3a-RMI1/2^, Mph1^FANCM^, and Srs2), imparts robustness in donor selection and contributes to the high fidelity of DSB repair by HR, which can otherwise lead to repeat-mediated chromosomal rearrangements (Putnam *et al*, 2009, 2016; Piazza & Heyer, 2019; Danilowicz *et al*, 2017).

Intuitively, transcription at the donor site appears incompatible with the co-occurrence of a D-loop, but priority rules between the two have largely remained unexplored. This is mainly due to (i) technical limitations in detecting D-loops in cells, and (ii) difficulties in disentangling the role of transcription in generating recombinogenic lesions from that of regulating their repair. Evidence in budding yeast and human hint at an inhibitory role of transcription at synaptic or post-synaptic HR steps. First, transcriptional activity biases the repair outcome of meiotic inter-homolog recombination towards non-crossovers in humans (McVicker & Green, 2010; Pouyet *et al*, 2017; Palsson *et al*, 2025). In budding yeast, early work quantifying spontaneous recombination rates between hetero-alleles with varying transcriptional levels also suggested that besides generating recombinogenic lesions transcription could also inhibit their repair (Saxe *et al*, 2000). More recently, initiation of break-induced replication (BIR) was shown to be impaired by highly-transcribed RNA Polymerase II (hereafter RNA PolII) genes present in head-on orientation in budding yeast (Liu *et al*, 2021; Uribe-Calvillo *et al*, 2022). BIR is a non-canonical, conservative and unstable Rad51-dependent replication process that takes place in the context of a D-loop structure, at least in its initial stages (Davis & Symington, 2004; Lydeard *et al*, 2007; Smith *et al*, 2007; Wilson *et al*, 2013; Mayle *et al*, 2015; Donnianni & Symington, 2013; Saini *et al*, 2013; Donnianni *et al*, 2019; Liu *et al*, 2021). It suggested a dominance of transcription over recombination, at least in the initial elongation step specific to this low-fidelity HR sub-pathway. However, how transcription might impinge on the core synaptic steps of canonical HR (*i.e.* on donor invasion and D-loop metabolism) remains unknown.

Here, using a site-specific DSB induction system and molecular assays for D-loop detection in *S. cerevisiae*, we show that donor transcription by RNA PolII suppresses D-loops in an orientation-dependent manner. We delineate its requirement and relationship with previously characterized D-loop disruption activities, and demonstrate its involvement in suppressing repeat-mediated rearrangements. These findings reveal the prioritization between two universal DNA-dependent processes and its function in genome maintenance.

## Results

### Experimental system

We used a well-established experimental system in haploid *S. cerevisiae* cells in which a site-specific DSB can be rapidly induced upon over-expression of the *HO* gene (Piazza *et al*, 2018, 2019). The HO cut-site (HOcs) at the *URA3* locus on chr. V is flanked on the left by a region of homology to a “donor” site present at the *LYS2* locus on chr. II (**Fig. 1A**). The right side has no donor, which purposefully precludes downstream repair steps. Absolute levels of D-loop joint molecules formed by the left DSB end at, and extended from this donor can be quantified over a ∼8 hours period using the proximity ligation-based D-loop-Capture (DLC) and D-loop Extension (DLE) assays, respectively (Piazza *et al*, 2018, 2019; Reitz *et al*, 2022). DLC uniquely requires *in vivo* inter-strand DNA crosslinking with psoralen. We used an improved DLC protocol in which psoralen was de-crosslinked prior to qPCR, enabling absolute determination of D-loop joint molecules (Reitz *et al*, 2022). In this system, >90% of DNA molecules are cut within 1 hour of *HO* expression induction, D-loops are first detected at 2 hours, peak at 3-4 hours, and are extended from 4 to 8 hours post-DSB induction (Piazza *et al*, 2019). Scoring D-loops at 2 hours thus focuses on D-loops prior to their extension.

**Figure 1:**
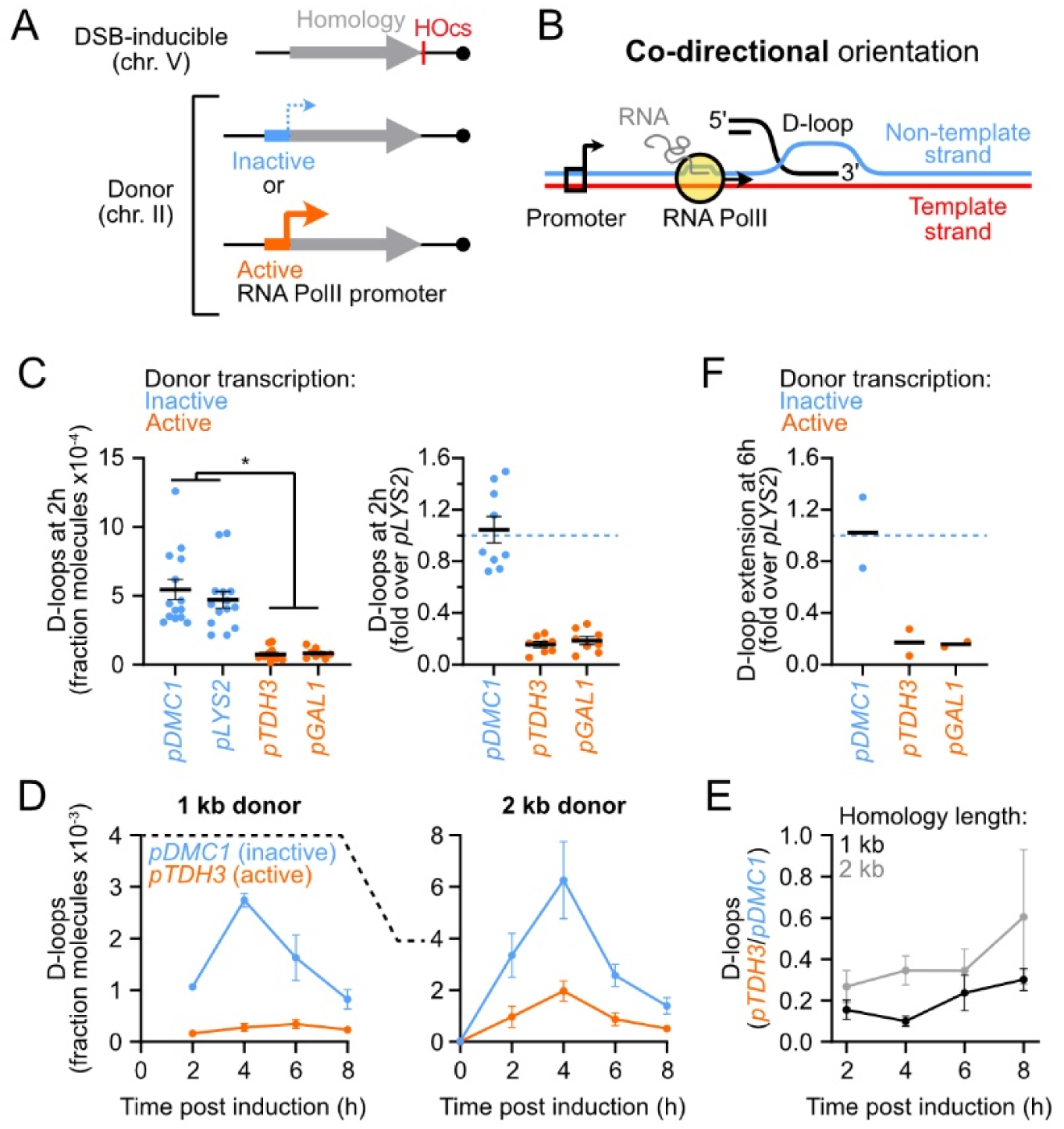
Co-directional donor transcription suppresses nascent D-loops. A) Experimental system with the donor transcribed in the co-directional orientation. B) Depiction of RNA PolII relative to the D-loop in the co-directional orientation. C) Left: Absolute D-loops detected 2 hours post-DSB induction with transcriptionally inactive (*pDMC1* and *pLYS2*) and active (*pTDH3* and *pGAL1*) donors (strains APY502, APY867, APY725, and APY724, respectively). Right: D-loop levels relative to that of a strain bearing a transcriptionally inactive *pLYS2* donor scored in a parallel. D) Absolute D-loops detected following DSB induction with two homology length at transcriptionally inactive and active donors (strains APY502, APY867, APY354, and APY1180). E) D-loop inhibition upon donor transcription, determined from data in E). F) D-loop extension at 6 hours post-DSB induction, normalized to that of a strain bearing a transcriptionally inactive *pLYS2* donor scored in a parallel. Strains are the same as in C). C-F) Data show individual biological replicates and mean ± SEM. * denotes statistical significance (p<0.05).

### Transcription of the donor reduces nascent D-loop levels

In order to investigate the effect of the RNA PolII-dependent transcriptional activity at the donor site on D-loop metabolism, we placed the 1 kb-long “L” donor present on chr. II under the control of promoters that drive either no or minimal (*pDMC1* and *pLYS2*) or high (*pTDH3* and *pGAL1*) donor transcription in our culture conditions (**Fig. 1A**). The promoter is not part of the homology region, and is oriented so that the same DNA strand is a template for transcription and recombination, which we refer to as the “co-directional” orientation (**Fig. 1B**). The expected transcriptional activity was verified by ChIP-qPCR of RNA PolII and RT-qPCR with these different constructs (**Fig. S1A-B**). Donor transcription did not affect the expression of the downstream gene *RAD16* either (**Fig. S1B**), and had no indirect effect on DSB formation on chr. V (**Fig. S1C**).

Strikingly, D-loop levels detected 2 hours post-DSB induction were ∼10-fold lower when the donor was transcribed than when it was not (**Fig. 1C**). The D-loop level was inversely correlated with the donor transcription level (**Fig. S1D**). This inhibition was observed at all time points up to 8 hours post-DSB induction (**Fig. 1D-E**), indicating that the inhibition observed at 2 hours was not due to a delay in D-loop formation. Similar results were obtained with an independent construct bearing 2 kb of homology (**Fig. 1D-E** and **S1E**). Consistently, the downstream step of D-loop extension scored 6 hours post-DSB induction was inhibited ∼7-fold upon donor transcription (**Fig. 1F**). These observations indicate that transcription of the donor interferes with the synaptic steps of recombination, by preventing D-loop formation and/or causing their disruption.

### Transcription suppresses D-loops acutely

In order to gain insights into the mechanism of transcription-dependent D-loop suppression, we sought to determine how quickly D-loops respond to transcriptional changes at the donor. To this end, we placed the donor under the control of a copper-inducible *pCUP1* promoter (Labbé & Thiele, 1999). D-loops were left to form in the absence of copper for ∼110 minutes post-DSB induction, and transcription was triggered upon copper addition ∼5-10 minutes prior to DNA crosslinking and D-loop detection (**Fig. 2A** and **S2A**). This short transcriptional induction was sufficient to cause a dose-dependent ∼2.5 to 11-fold drop in D-loop levels (**Fig. 2A**). D-loops formed downstream of the silent *pDMC1* promoter were not affected by copper addition, ruling out indirect effect of copper on D-loop metabolism (**Fig. 2A**).

**Figure 2:**
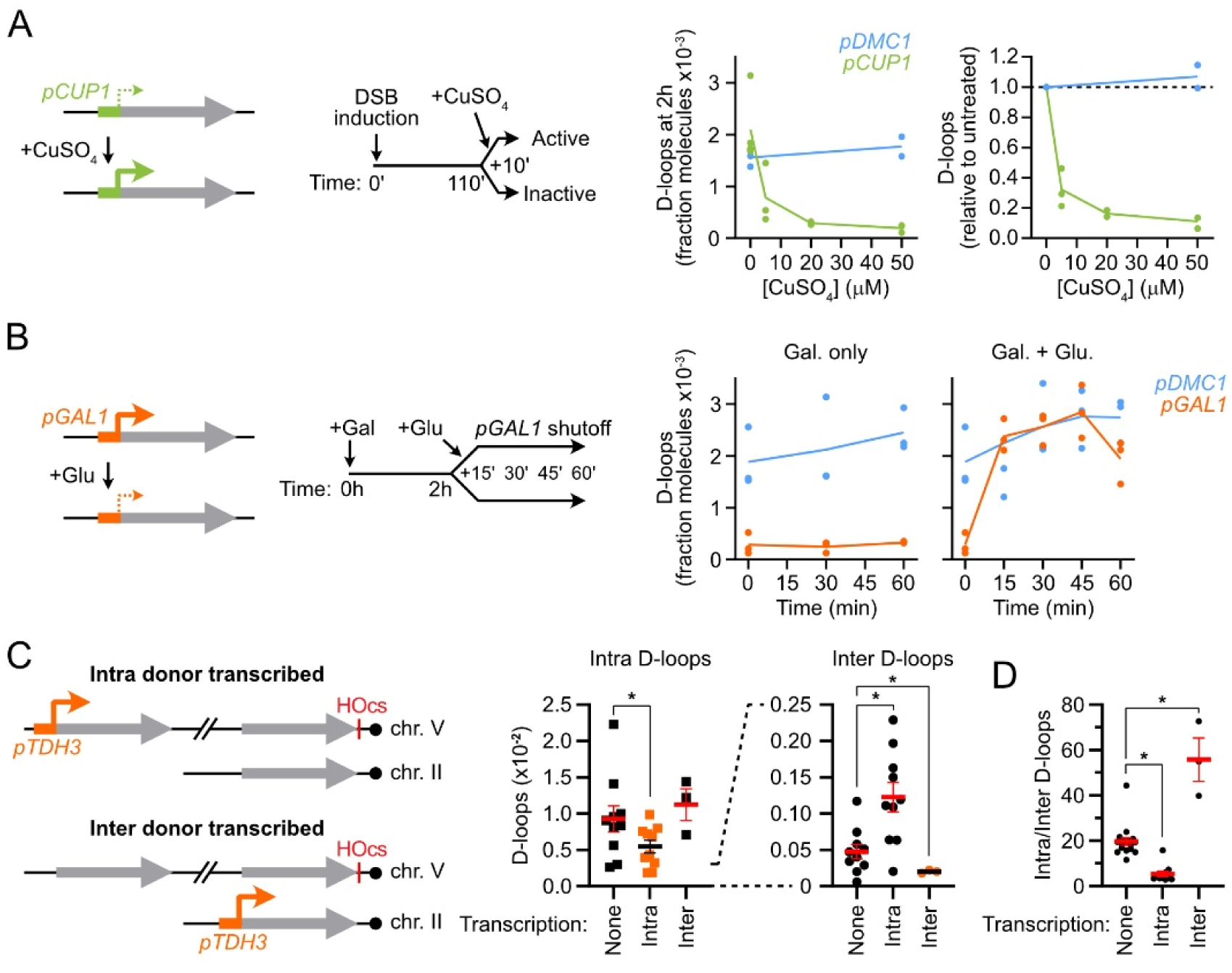
Donor transcription suppresses D-loops in *cis*. A) Copper-induced transcriptional activation at the donor from the *pCUP1* promoter ∼10 minutes prior to D-loop detection causes a dose-dependent D-loop loss (APY503). The control *pDMC1* promoter is not copper-responsive (APY502). B) Kinetics of D-loop restoration following transcriptional shutdown at the donor from the *pGAL1* promoter upon glucose addition (APY725). The control *pDMC1* promoter is not sensitive to the carbon source (APY502). C) Donor transcription inhibits D-loop formation in *cis*. Absolute D-loops formed at the intra donor (left) and the inter donor (right) in the absence of donor transcription (APY809), upon transcription of the intra donor (APY1587), or upon transcription of the inter donor (APY1709). D) Fold preference for D-loop formation at the intra over the inter donor. From data in C). A-D) Data show individual biological replicates (points) and mean ± SEM. * denotes statistical significance (p<0.05).

Conversely, we determined the kinetics of D-loop recovery following the glucose-induced shut-off of transcription at a *pGAL1* promoter (Nehlin *et al*, 1991) (**Fig. 2B**). D-loops levels are recovered within 15 minutes post-glucose addition and remained equivalent to the non-transcribed *pDMC1* donor afterwards (**Fig. 2B** and **S2B**).

These rapid responses to transcriptional activation and shutoff suggest that transcription directly undermines the stability of D-loops (or of the precursor synaptic complex) in *cis*. In the following sections, we examine this possibility by investigating the role of secondary consequences of transcriptional activity on D-loop levels.

### Transcription suppresses D-loops in *cis*

To directly address whether transcription of the donor exerts its effect in *cis*, or whether it acts in *trans* through its RNA product, we determined D-loop formation at the level of two competing donors, only one of which being transcribed. If the transcription exerts its inhibitory effect in *trans*, D-loop levels should be reduced at both donors. If transcription acts in *cis*, D-loop levels should only be reduced at the transcribed donor, and may even redirect D-loop formation onto the other, non-transcribed donor it competes with.

We used a previously characterized system with one donor located on the same chromosome as the DSB site (intra donor) and one located on chr. II (inter donor) (Piazza *et al*, 2021) (**Fig. S2C**). We confirmed with our improved DLC protocol (Reitz *et al*, 2022) that D-loops formed with a ∼20-fold preference at the intra over the inter donor in the absence of transcription (Piazza *et al*, 2021) (**Fig. S2C-D**). Presence or absence of an inter donor did not detectably affect intra D-loop formation (**Fig. S2C**). On the contrary, presence of the intra donor caused a ∼4- to ∼20-fold reduction in inter D-loops (**Fig. S2E**). Hence, presence of the intra donor outcompetes the inter donor for D-loop formation.

Transcription of the intra donor caused a significant 1.8-fold decrease of intra D-loops, and led to a 2.6-fold increase of inter D-loops (**Fig. 2C**). It resulted in a 3.7-fold decrease in the intra/inter donor preference (**Fig. 2D**). Conversely, transcription of the inter donor caused a significant 2.4-fold decrease of inter D-loops without detectably affecting intra D-loops, which led to a 2.8-fold increase in the intra/inter donor preference (**Fig. 2C-D**). These results establish that transcription specifically reduces D-loop levels at the transcribed donor, demonstrating that transcription suppresses D-loops in *cis.* Furthermore, this *cis*-acting inhibition can redirect D-loop formation onto the non-transcribed donor, swinging the intra/inter donor preference up to 10-fold (**Fig. 2D**).

### Transcription suppresses D-loops independently of RNA:DNA hybrids

RNA:DNA hybrids-containing three-stranded structures called R-loops may form co-transcriptionally in *cis* at certain highly-transcribed genomic loci (Gómez-González & Aguilera, 2021). Low-level RNA:DNA hybrids could be detected at the locus used as a donor in our system (*i.e. LYS2*) by H-CRAC in mutant contexts (Aiello *et al*, 2022) or by DRIP-qPCR when it was artificially over-expressed (Mérida-Cerro *et al*, 2024).

In order to addressed whether RNA:DNA hybrids were involved in transcription-dependent D-loop inhibition, we evaluated D-loop levels in various contexts reported either (i) to eliminate RNA:DNA hybrids upon over-expression of RNAseH1 (Wahba *et al*, 2011), or (ii) to exacerbate their formation and/or stability, in mutants of the THO complex (*mft1*Δ) or of the transcription elongation factor TFIIS (Huertas & Aguilera, 2003; San Martin-Alonso *et al*, 2021). RNAseH1 was over-expressed from a *pGAL1* promoter on a multi-copy 2µ plasmid together with DSB induction upon galactose addition. RNAseH1 over-expression did not significantly affect D-loop levels, neither with the *pDMC1* nor the *pTDH3* constructs (**Fig. 3A** and **S3A**). Likewise, the THO complex *mft1*Δ mutant and the *tfiis*Δ mutant did not cause a reduction in D-loop levels (**Fig. 3B**). On the contrary, the *tfiis*Δ mutant partly relieved the transcription-dependent D-loop inhibition (**Fig. 3B**). These observations indicate that RNA:DNA hybrids are not involved in suppressing D-loops at highly transcribed genes. Given the role of TFIIS in promoting transcription elongation by RNA PolII (Sigurdsson *et al*, 2002; Zatreanu *et al*, 2019), it further suggests that D-loop suppression requires efficient transcription across the donor.

**Figure 3:**
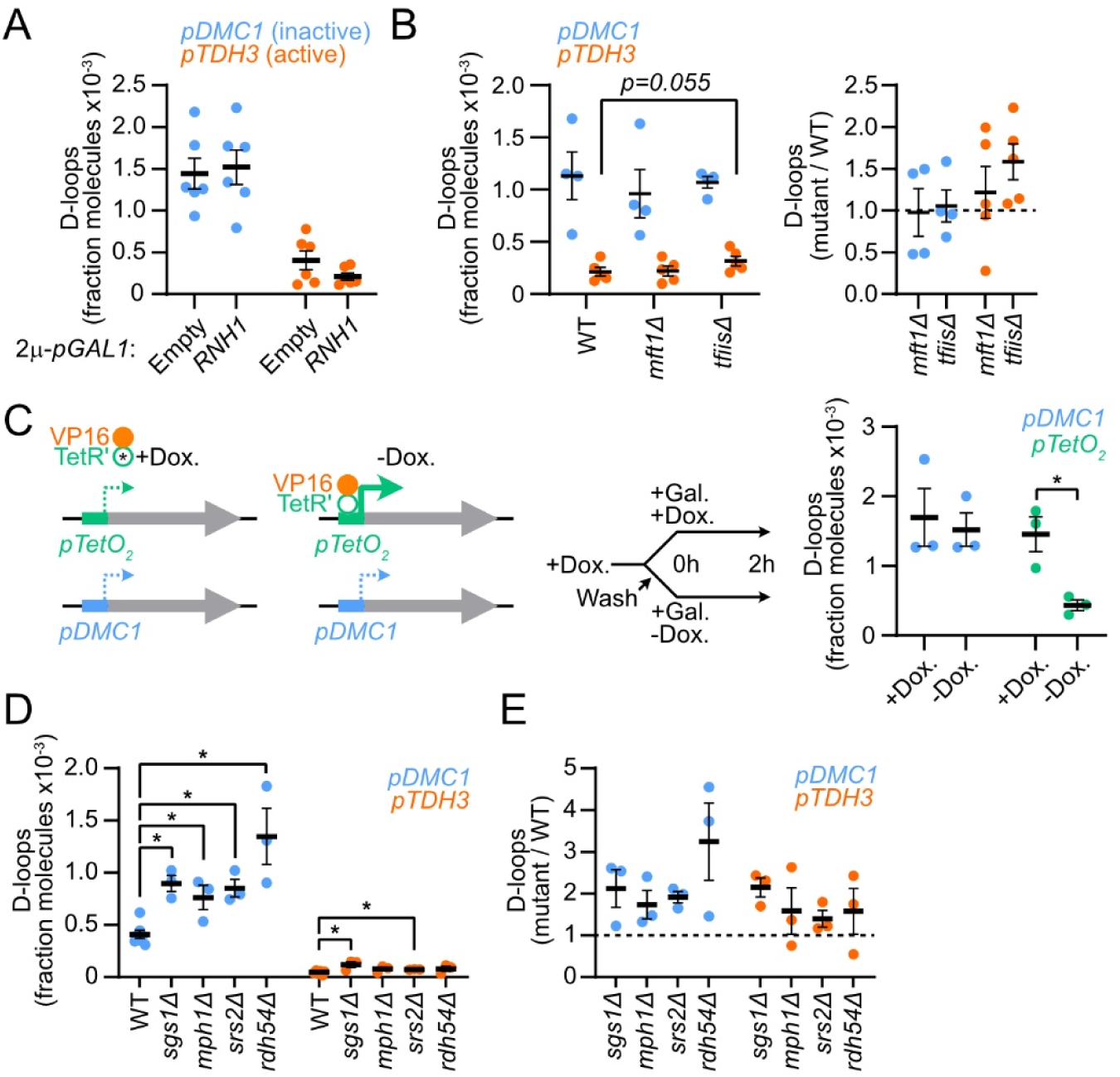
Genetic determinants of transcription-mediated D-loop suppression. A) Absolute D-loop levels are unaffected by RNAseH1 over-expression, irrespective of the transcriptional status of the donor (APY1272, APY1274, APY1278, and APY1280). B) Left: absolute D-loop levels at transcriptionally inactive and active donors in WT (APY502 and APY725), *mft1*Δ (APY1114 and APY1116) and *tfiis*Δ (APY1485 and APY1487) strains. Right: relative mutant values compared to a WT strain assayed in parallel. C) Left: Rationale and experimental scheme for transcriptional activation by a heterologous TetR-VP16 construct. Right: Absolute D-loop levels (APY1292 and APY1294). D) Left: absolute D-loop levels at transcriptionally inactive and active donors in WT (APY502 and APY725), *sgs1*Δ (APY824 and APY795), *mph1*Δ (APY789 and APY796), *srs2*Δ (APY791 and APY798), and *rdh54*Δ (APY793 and APY799) strains. Right: relative mutant values compared to a WT strain assayed in parallel. A-D) Data show individual biological replicates (points) and mean ± SEM. * denotes statistical significance (p<0.05).

### Transcription suppresses D-loops independently of peripheral nuclear delocalization and endogenous transcription initiation factors

Transcriptional activation can lead to the delocalization of the locus from the nuclear lumen to the nuclear periphery (Casolari *et al*, 2004; Brickner & Walter, 2004). However such delocalization has been reported to occur ∼15 minutes post-transcriptional induction at the earliest (Randise-Hinchliff *et al*, 2016), while less than 10 minutes of transcriptional induction was sufficient to cause full D-loop disruption (**Fig. 2A**). Furthermore, *pGAL1* retains its peripheral nuclear localization for >14 hours post-transcriptional shutoff with glucose (Sood *et al*, 2017), yet D-loops we recovered in less than 15 minutes in these conditions (**Fig. 2B**). These results indicate that the peripheral nuclear localization of highly transcribed genes is not implicated in D-loop suppression.

In order to confirm this independence, and address the role of endogenous transcription initiation factors in mediating D-loop suppression, we placed the donor under the control of a heterologous doxycycline-responsive dual activator/repressor system (Bellí *et al*, 1998). In the absence of doxycycline, the TetR’ DNA binding domain of *E. coli* fused to the transcription activator VP16 from the SV40 virus drives transcription from a *pTetO_2_* promoter, placed upstream of the donor (**Fig. 3C**), while the TetR-Ssn6 fusion suppresses transcription otherwise. This VP16 fusion strongly stimulates transcription in budding yeast while retaining the transcribed locus in the nuclear lumen (Sadowski *et al*, 1988; Garí *et al*, 1997; Taddei *et al*, 2006). Cells grown in the presence of doxycycline (*i.e.* in which TetR’-VP16 does not associate to its *TetO* target) exhibited similar D-loop levels whether the donor was under the control of the *pTetO_2_* or the control *pDMC1* promoter (**Fig. 3C**). Transcriptional activation from *pTetO_2_* promoter upon doxycycline removal caused a 3.5-fold D-loop loss (**Fig. 3C**). D-loop levels remained unaffected with the *pDMC1* promoter, ruling out indirect effects of doxycycline or the TetR-VP16 construct on early recombination steps. These results confirm that transcription-mediated D-loop inhibition occurs independently of the delocalization of the transcribed locus at the nuclear periphery. It also shows that such inhibition can be triggered by a heterologous transcription initiation factor, and is thus independent on any specific endogenous transcription initiation factor.

### Transcription suppresses D-loops independently of STR, Srs2, Mph1 and Rdh54

The Sgs1-Top3-Rmi1^BLM-TOPO3a-RMI1/2^ (STR) helicase-topoisomerase complex, the Mph1^FANCM^ and Srs2 helicases, and the Rdh54^RAD54B^ dsDNA translocase suppress HR- and repeat-mediated gross chromosomal rearrangements (Putnam *et al*, 2009, 2016), disrupt or alter formation of D-loops in reconstituted *in vitro* reactions with Rad51 and Rad54 (Prakash *et al*, 2009; Fasching *et al*, 2015; Liu *et al*, 2017; Shah *et al*, 2020), and cause a reduction of the amount of nascent D-loops detected in our system (Piazza *et al*, 2019; Xie *et al*, 2024). To address the genetic interactions between transcription and these *trans* D-loop disruption factors, we combined these individual mutations with a donor under the control of the silent *pDMC1* or the active *pTDH3* promoter. The 2- to 3-fold increase in D-loop levels detected in these mutants with the non-transcribed donor recapitulated previous findings obtained with the donors under the endogenous *pLYS2* promoter (Piazza *et al*, 2019) (**Fig. 3D-E**). None of them relieved the ∼10-fold inhibition imposed by transcription (**Fig. 3D**), indicating that transcription-mediated D-loop suppression does not require any of these factors. Moreover, the fold-change in these mutants over the wild-type background was overall similar with the silent and active promoters (**Fig. 3E**). Consequently, donor transcription is a distinct D-loop suppression pathway without detectable overlap with that conferred by these specialized HR regulators. Finally, the absolute fold-change in D-loop levels measured in these mutants (2- to 3-fold) vs. that conferred by transcription (up to ∼10-fold) shows that transcription can be the main D-loop suppression pathway in cells, as a function of the local transcriptional activity.

The mismatch repair protein Msh2, also involved in heteroduplex rejection, was not implicated in D-loop reversal at our perfectly homologous substrates, irrespective of the transcriptional status of the donor (**Fig. S3B**).

### The efficiency of D-loop suppression depends on transcription directionality

We addressed the impact of the directionality of transcription relative to that of the D-loop in two contexts: by repositioning the promoter at the donor site; or by inverting the region of homology near the break site (**Fig. 4A**). In these two independent “head-on” contexts, the RNA PolII is set to encounter the 3’ junction of the D-loop rather than the 5’ junction (**Fig. 4A**).

**Figure 4:**
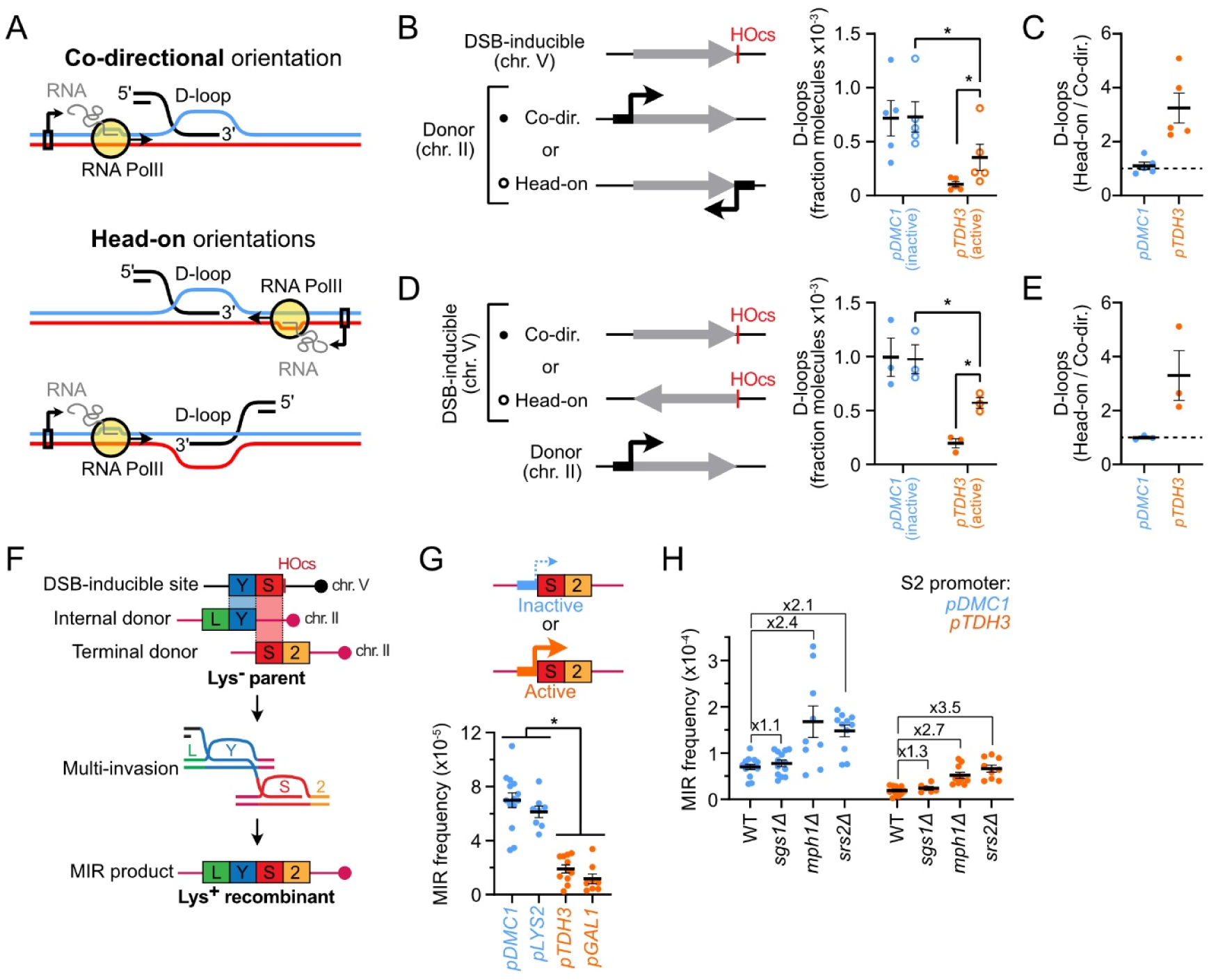
Effect of transcription directionality and role of transcription in suppressing repeat-mediated chromosomal rearrangements. A) Depiction of RNA PolII relative to the D-loop in the co-directional and head-on orientations. B) Inverting the transcription directionality at the donor by repositioning the promoter partly rescues D-loop levels when transcription is active (APY502, APY941, APY725, and APY999). C) Fold difference in D-loop levels in the head-on compared to the co-directional orientation. Data from B). D) Inverting the transcription directionality at the donor by inverting the homology sequence near the DSB partly rescues D-loop levels when transcription is active (APY502, APY1731, APY725, and APY1733). E) Same as C), with data from D). F) Tripartite inter-chromosomal genetic system to study MIR. G) Co-directional transcription of the terminal donor suppresses MIR (APY1130, APY1512, APY1077 and APY1075). H) MIR with a transcriptionally inactive and active terminal donor in WT (APY1130 and APY1077), *sgs1*Δ (APY1693 and APY1692), *mph1*Δ (APY1682 and APY1681), and *srs2*Δ (APY1684 and APY1683) strains. B-E, G-H) Data show individual biological replicates (points) and mean ± SEM. * denotes statistical significance (p<0.05).

Transcriptional activity at the donor was similar with promoters placed in the co-directional and head-on orientations, as verified by ChIP-qPCR against the Rpb1 subunit of RNA PolII (**Fig. S4A**). Donor transcription in the head-on orientation led to reduced D-loop levels, but the extent of this inhibition was less pronounced compared to the co-directional orientation (**Fig. 4B-C**). Specifically, D-loop levels were ∼3-fold higher at donors transcribed in the head-on compared to the co-directional orientation (**Fig. 4C**). D-loops at the non-transcribed donor were not affected by promoter repositioning (**Fig. 4B-C**). Likewise, inversion of the homology region on the broken molecule partly alleviated the inhibition posed by donor transcription, while exerting no effect on D-loop levels at the non-transcribed donor (**Fig. 4D-E**). Again, D-loop levels were ∼3-fold higher with transcription in the head-on than in the co-directional orientation (**Fig. 4E**). These two independent sets of constructs show that transcription is not as effective at causing D-loop loss in the head-on than in the co-directional orientation. Consequently, a permissive orientation exists for D-loops to form, and thus for HR to proceed from one DSB end, in highly transcribed gene.

### Donor transcription protects against chromosomal rearrangements

Rad51-ssDNA filaments can form multiple D-loops along their length *in vitro* and in *S. cerevisiae* cells, resulting in multi-invasions (MI) DNA joint molecules (Wright & Heyer, 2014; Piazza *et al*, 2017) (**Fig. 4F**). Endonucleolytic processing of these HR byproducts are at risk of translocating the intact donors engaged in adjacent D-loops in a general process refer to as multi-invasion recombination (MIR) (Piazza *et al*, 2017; Reitz *et al*, 2023). In order to address whether transcription-dependent D-loop suppression could protect against MIR, we adapted a previously established genetic assay in which the DSB-inducible site, the internal donor, and the terminal donor are on different chromosomes (Piazza *et al*, 2017) (**Fig. 4F**). We placed the terminal donor under the control of inactive or low activity promoters (*pDMC1, pLYS2*) or highly active promoters (*pTDH3, pGAL1*) in our culture conditions (**Fig. 4G**). The internal promoter remained under the control of its native *pLYS2* promoter for selection purposes. Transcription of the terminal donor caused a 3.2- to up to 6-fold decrease in MIR frequency (**Fig. 4G**). In contrast, cells deficient for Sgs1, Mph1 and Srs2 caused MIR to increase by only ∼1.2 to 3-fold in the absence of transcription (**Fig. 4H**). Hence, transcription of a single donor inhibits MIR more effectively than any of these *trans* HR regulators. These mutants caused similar fold increases in MIR whether or not the terminal donor was transcribed, corroborating genetically the independence between transcription and these HR regulators in promoting D-loop disruption determined molecularly (**Fig. 3D-E**). In conclusion, transcription inhibits the formation of HR-mediated chromosomal rearrangements with efficiency that can exceed that conferred by conserved HR regulators involved in promoting genome maintenance.

## Discussion

Here, through direct D-loop detection in multiple genetic and mutant contexts, we show that donor transcription by RNA PolII suppresses D-loops (and/or its precursor paranemic Rad51-bound synaptic complex). Functionally, this layer of HR regulation promotes genome maintenance by inhibiting repeat-mediated chromosomal rearrangements. Hence, homologous recombination fidelity is not equally enforced along the genome, and instead depends on the local transcriptional activity.

### Putative mechanism of transcription-mediated D-loop suppression

D-loop suppression at a highly-transcribed donor occurs in *cis* and can be rapidly (<10 min) switched on and off, strongly suggesting that D-loop suppression is a direct consequence of RNA PolII translocation at the donor. Supporting this suggestion, we could rule out secondary consequences of transcription such as peripheral nuclear delocalization, RNA:DNA hybrids, and the RNA product acting in *trans* in D-loop suppression. It sets this mechanism of HR control apart from those implicating RNA molecules identified in yeast and human cell lines (Keskin *et al*, 2014; Meers *et al*, 2020; Ouyang *et al*, 2021; Liu *et al*, 2022).

RNA PolII is a processive directional molecular motor threading along dsDNA, a process facilitated by TFIIS (Gnatt *et al*, 2001; Kettenberger *et al*, 2004; Charlet-Berguerand *et al*, 2006). Such active translocation may dissociate already formed D-loops by mechanically migrating its strand exchange junctions, an energetically neutral reaction. The fact that transcription mainly causes loss of co-directional D-loops (i) suggests that it does not solely act by preventing the upstream step of Rad51-ssDNA NPF binding to dsDNA at that site, and (ii) makes it unlikely that D-loop dissociation is primarily mediated by topological changes at the donor. It instead suggests that the prioritization of transcription over the synaptic steps of HR depends on the type of DNA strand exchange junctions encountered by RNA PolII, and/or of the presence of HR proteins decorating them relative to an incoming RNA PolII. Consistently, RNA PolII obtained from HeLa cells extract could traverse a model co-linear D-loop substrate *in vitro*, while a head-on D-loop was a roadblock (Pipathsouk *et al*, 2017). The precise structural basis for this orientation-dependent behavior, at the heart of the prioritization between the two processes, remains to be determined.

### Role of transcription-recombination prioritization in genome maintenance

Transcription has primarily been recognized has a pro-recombinogenic process through the formation of replication-dependent DNA lesions. Here we describe a downstream function of transcription that mitigates the detrimental consequences of these lesions on genome maintenance by impinging on the recombination process. It builds upon the earlier proposal of an anti-recombination function of transcription by the Jinks-Robertson laboratory (Saxe *et al*, 2000). We precise the nature and extent of this regulation by showing that a permissive orientation for D-loop formation likely ensures that HR repair of DSBs formed at highly-transcribed genes can occur. Instead of an anti-recombination mechanism, we propose that transcription locally increases the stringency of the donor selection process by requiring several rounds of DNA strand invasion prior to D-loop extension; a kinetic proof-reading strategy likely to disproportionately disfavor usage of short, homeologous, and/or distant repeated sequences (Piazza & Heyer, 2019). Consistently, we noted that long homologies (**Fig. 1E**) and DSB-donor spatial proximity (**Fig. 2C**), two hallmarks of the sister chromatid, partly counteracted the effect of transcription, presumably by promoting rapid re-invasion at the same locus. Finally, the inhibition of DNA strand invasion by both DSB ends is likely to disfavor the formation of double Holliday Junction intermediates, and thus suppress the formation of crossover repair products. This mechanism may underlie the observed bias towards the non-crossover repair outcome at transcribed genes in human meiosis (McVicker & Green, 2010; Pouyet *et al*, 2017; Palsson *et al*, 2025).

## Acknowledgements

We thank the Piazza, Bernard, and Jost lab members, as well as Wolf-Dietrich Heyer, Bertrand Llorente and Laurent Duret for stimulating discussions and helpful suggestions. We are grateful to Vincent Vanoosthuyse for stimulating discussions and its critical reading of the manuscript and Lorraine Symington for sharing plasmids and strains.

This research was supported by the European Research Council (ERC) under the European Union’s Horizon 2020 to AP (ERC grant agreement 851006) and a 4^th^ year PhD fellowship extension from the Fondation pour la Recherche Médicale to YD (FDT202404018451).

## Author contributions

Conceptualization: AP; Experiments: YD, AP; Data analysis: YD, AP; Data interpretation: YD, AP; Supervision: AP; Funding Acquisition: YD, AP; Writing: YD, AP.

## Declaration of interest

None

## STAR Methods

### Resources availability

#### Lead contact

Further information and requests for resources and reagents should be directed to and will be fulfilled by the lead contact, Aurèle Piazza.

#### Materials availability

*S. cerevisiae* strains generated in this study are available upon request. The strains genotype is listed in **Table S1**.

### Experimental Model and Study Participant Details

#### Saccharomyces cerevisiae strains

The haploid and diploid *Saccharomyces cerevisiae* strains used in this study derive from the W303 *RAD5^+^* background. The strains genotype is provided in **Table S1**. The annotated sequences of all the DSB-inducible and donor constructs are provided in **Dataset S1**.

The *HO* gene was placed under the control of the *pGAL1*/*10* promoter at *TRP1* locus on chromosome IV (Pannunzio *et al*, 2008), the point mutation at the HO cut-site (*HOcs*) present at the mating-type locus (*MAT*) on chromosome III to prevent its cleavage by HO (*MAT*a-inc and *MAT*α-inc) and the DSB-inducible construct have been described previously (Piazza *et al*, 2019, 2018; Reitz *et al*, 2022). Briefly, The DSB-inducible construct consists of the *HOcs* introduced in place of the *URA3* gene on chromosome V (-16 to +855 from the start codon). The *HOcs* is flanked on its left side by a 1 or 2 kb-long fragment of the 5’ end of the *LYS2* gene minus the start codon, and are referred to as “*L*” and “*LY*”, respectively. The “*L*” and “*LY*” donors are located at the endogenous *LYS2* locus on chr. II (the “*YS2*” or “*S2*” remainder of the gene has been removed) and is referred to as the “donor”. The lack of homology between the donor and the right side of the break prevents repair by synthesis-dependent strand annealing or DSBR (Pâques & Haber, 1999; Szostak *et al*, 1983). The orientation of the donor towards *CEN2* prevents repair completion by BIR (Morrow *et al*, 1997; Pham *et al*, 2021). This system thus precludes repair completion, so as to retain a constant number of cells undergoing repair at all time points (Reitz *et al*, 2022).

In the *L*-inverted construct, the orientation of the “*L*” sequence is flipped but remains on the left side of the *HOcs*. In the co-directional orientation, these donors are under the control of the native *pLYS2* promoter (-155 to -1 bp from the TSS), the *pDMC1*, the *pTDH3*, the *pGAL1,* or the *pCUP1* promoters. The *pTetO_2_-TATA* promoter was obtained from the pST1873 vector. The S288c coordinates of the promoters are listed in **Table S2**. In the head-on orientation, the promoters and the *tLYS2* terminator (+1 to +70 bp from the stop codon) were exchanged and reverse-complemented. The artificial *tGuo1* terminator (5’-TATATAACTGTCTAGAAATAAAGGTGCAGGCATTTCAAA-3’) (Curran *et al*, 2015) was further added downstream of *tLYS2*.

Strains with a second donor, which is inserted 85 kb away from the DSB site at the *CAN1* locus on chromosome V, are used to compare the intra-chromosomal and inter-chromosomal donors.

The diploid strains for MIR study were as in (Piazza *et al*, 2017). They contain a heterozygous DSB-inducible construct at *ura3* on chromosome V. The DSB-inducible construct consists of the *HOcs* flanked by the central, 2 kb-long part of the *LYS2* gene (YS). The donor in these strains is divided into two parts on each copy of chromosome II; the first copy carries the first half of the *LYS2* gene (*LY*), and the second copy carries the second half of the *LYS2* gene (*S2*), without sequence overlap.

The *RNAseH1* (*RNH1*) was over-expressed from the *pGAL1* promoter on a the 2µ plasmid pYES2 (Invitrogen cat. V82520). The empty pYES2 plasmid was used as a negative control.

#### Culture media

Culture media were prepared according to standard protocols (Treco & Lundblad, 2001). Yeast Extract Peptone media (YP) are composed of 1% bacterial and yeast extract and 2% peptone, with variations in the carbon source: 2% dextrose (YPD), 2% lactate (YPL), or 2% galactose (YPGal). The synthetic medium contained 0.17% yeast nitrogen base, 0.5% ammonium sulfate, 2% dextrose, and 0.2% all amino acids for the synthetic complete (SC) or 0.2% appropriate amino acids dropout for the synthetic selective media (SD-AA). The synthetic dropout galactose contained 2% galactose. Additionally, the solid media contained 2% agarose.

Note that the water used in media preparation has changed over the course of this study from osmosis water to ultrapure water obtained after Milli-Q filtering, which improved yeast growth and led to slight changes in absolute D-loop levels. Consequently, D-loop levels were always compared between matched samples grown in the same media and processed in parallel.

#### Induction of DSB and cell treatments

*HO* expression was induced upon addition of 2% galactose to YPL cultures reaching OD600 ∼0.5.

In the case of *pCUP1* activation with copper, cultures were grown in SC-Lactate media up to OD600 ∼0.5 and *HO* expression induced upon addition of 2% galactose. Copper sulfate salt (CuSO_4_ in water) was added at 5, 20 or 50 µM 110 minutes post-DSB induction, corresponding to 10’ prior to DLC sample collection.

In the case of RNAseH1 over-expression and its matched empty vector control, saturated precultures prepared in 2% glucose-containing SD-Uracil media were diluted in lactate-containing synthetic dropout lacking uracil and grown overnight to OD600 ∼0.5. *HO±RNH1* overexpression was induced upon addition of 2% galactose.

In the case of transcriptional control of the donor by TetR-Ssn6 and TetR-VP16, exponential cultures in SC-Lactate reaching OD600 ∼0.5 were split and 20 µg/mL doxycycline (Sigma-Aldrich cat. D9891) or the equivalent ethanol concentration was added together with 2% galactose.

#### D-loop Capture (DLC) assay

The DLC was performed as in (Piazza *et al*, 2019; Reitz *et al*, 2022), including the psoralen crosslink reversal step described in (Reitz *et al*, 2022). Briefly, a site-specific DSB was induced upon over-expression of the HO endonuclease by adding galactose at a final concentration of 2% to an exponentially growing cell culture in YEP-lactate media. 2.10^8^ cells we collected prior to, and at various time points after galactose addition. For DLC, cells were re-suspended in crosslinking solution (0.1 mg/mL Trioxsalen (Sigma-Aldrich T6137), 50 mM tris HCl pH 8.0, 50 mM EDTA, 20% ethanol) and the DNA was crosslinked with ∼32 J/cm^2^ UV-A (365 nm) irradiation in a Bio-link – BLX365 (Vilber-Lourmat, cat. 611110831) with permanent orbital agitation (∼50 rpm). Cell were spheroplasted for 15 min at 37°C with 3.5 µg/mL Zymolyase 100T in spheroplasting buffer (0.4 M sorbitol, 0.4 M KCl, 40 mM phosphate buffer pH 7.4, 0.5 mM MgCl_2_) and washed twice with spheroplasting buffer and three times with 1X Cutsmart buffer (20 mM tris acetate pH 7.9, 50 mM potassium acetate, 10 mM magnesium acetate, 100 µg/mL BSA) at 4°C. Pellets were resuspended in 1.4X Cutsmart buffer, flash frozen in liquid nitrogen and stored at -70°C. Cells were lysed upon addition of 0.1% SDS at 65°C for 10-15 min in the presence of a 80mer oligonucleotide (APO563; **Table S3**), whose annealing restores the EcoRI restriction site on the resected broken molecule. DNA was recovered from the spheroplasts, digested by EcoRI-HF (NEB, cat. R3101L), and ligated at low concentration (∼1.8 × 10^4^ genomes/µ l). DNA was extracted with phenol-chloroform after protein degradation using proteinase K. Psoralen inter-strand crosslinks and adducts was reversed in 100 mM KOH at 90°C for 30°C. The pH was neutralized upon addition of 66 mM of NaoAc pH 5.2. Approximately 6 × 10^5^ genome equivalent were used per quantitative PCR (qPCR) reaction, performed in duplicate, on a CFX96 Touch Deep Well Real-Time PCR Detection System (Bio-Rad cat. 3600037), using the SsoAdvanced Universal SYBR Green Supermix (Bio-Rad, cat. 1725274), following manufacturer’s instructions. Primers used are listed in **Table S3.** qPCR analysis were performed as described in (Reitz *et al*, 2022) using Bio-Rad CFX Maestro and Microsoft Excel.

#### D-loop extension (DLE) assay

The DLE was performed as in (Piazza *et al*, 2018; Reitz *et al*, 2022). Samples were collected and processed as for the DLC assay, except that no psoralen crosslinking was performed. At the lysis step, HindIII restrictions sites were restored upstream of the region of homology on the broken molecule and downstream of the donor site upon annealing of APO581 and APO640, respectively (**Table S3**). DNA was digested with HindIII-HF (NEB, cat. R3104L) instead of EcoRI. Quantitative PCR was performed using primers listed in **Table S3** and data analyzed as described in (Reitz *et al*, 2022).

#### RNA extraction and RT-qPCR

RNA extraction was performed using the Nucleospin RNA kit (Machery Nagel, ref: MN06 740588.250) following manufacturer’s instructions. The quantitative reverse transcription PCR (RT-qPCR) was performed on a CFX96 Real-Time System (Bio-Rad), using an iTaq™ Universal SYBR® Green One-Step Kit (Bio-Rad, cat. 1725274) following manufacturer’s instructions. Primers used are listed in **Table S3**.

#### Quantitative chromatin immunoprecipitation (ChIP-qPCR)

Approximately 1.5x10^8^ cells were crosslinked 2 hours post-DSB induction in 3% formaldehyde. Cell lysis was performed in 300 µ l of lysis solution (50 mM HEPES-KOH pH8, 140 mM NaCl, 1 mM EDTA pH8, 1% Triton X-100, 0.1% Sodium Deoxycholate, and 1 mM PMSF supplemented with a protease inhibitor cocktail (Roche cat. 11836170001)) with ø 500 µm acid-washed beads using a Precellys (6,800 rpm for 12 seconds, rest in ice for 45 seconds, repeated 4 times). The chromatin was sheared by sonication using a Covaris (240 W peak power, 20% duty factor and 200 cycles for 10 minutes). The lysate was clarified by centrifugation at 10,000 g at 4°C. RNA PolII immunoprecipitation was performed using a mouse anti-RNA polymerase II antibody (anti-Rpb1 clone CTD4H8, Sigma-Aldrich cat. 05-623) overnight followed by 2 hours incubation with equilibrated Dynabeads proteins G (Thermofisher cat. 10003) at 4°C. Beads were washed 3 times in lysis solution, 3 times in lysis solution supplemented with high salt (500 mM NaCl), twice with a wash solution (10 mM Tris-HCl, 500 mM LiCl, 1 mM EDTA pH 8, 0.5% Igepal CA-630, 0.1% Sodium Deoxycholate, 1 mM PMSF), and once with the TE-Na buffer (10 mM Tris-HCl pH 8, 1 mM EDTA, 50 mM NaCl). Elution was performed using 1% SDS at 65°C and the crosslink was reversed with 3% SDS overnight at 65°C. Following RNA degradation using RNase A and protein degradation using proteinase K, DNA was purified upon phenol-chloroform extraction and ethanol precipitation and resuspended in TE buffer (10 mM Tris-HCl pH 8, 1 mM EDTA). DNA was quantified using a Qubit dsDNA HS assay kit (Thermofisher cat. Q32854). RNA PolII occupancy of the donor was analyzed by quantitative PCR using primers targeting the 3’ and 5’ ends of the donor (**Table S3**). The IP over input ratio was normalized over that at the *ARG4* gene.

#### MIR translocation assay

The MIR translocation assay was performed using diploid *S. cerevisiae* as described in (Piazza et al., 2017), except that the DSB was not induced upon galactose addition in liquid media, but upon direct plating of an exponential culture in liquid YEP-lactate media on galactose-containing plates. Cells were spread on YPGal and YPD to determine viability, and on SDGal-Lysine and SDGlu-Lysine to determine the frequency of Lys+ recombinants in presence and absence of DSB, respectively. The viability and Lys^+^ frequencies were determined by counting colonies after incubating the plates for 2-3 days at 30°C.

### Quantification and statistical analysis

#### Statistical analysis

Most statistical comparisons were performed using a non-parametric two-tailed Mann-Whitney-Wilcoxon test. When n<4, a two-tailed Student t-test was used. Statistical tests were performed with GraphPad Prism 10. The statistical significance α cutoff was set at 0.05.

Each biological replicate is represented as an individual point in the figures. The nature of the summary representation (mean, SEM, etc.) of the data is indicated in each figure legend.

#### Software

Graphpad Prism 10

Microsoft Excel 2019

Bio-Rad CFX Maestro 1.1

### Supplemental Table Titles

Table S1: Genotype of the haploid *Saccharomyces cerevisiae* strains used in this study (related to STAR Methods).

Table S2: Genomic coordinates of the promoters used in this study (related to STAR Methods).

Table S3: Primers used in this study (related to STAR Methods).

### Supplemental item titles

**Data S1: Annotated sequences of the constructs used in this study (related to STAR Methods).**

## Notes

### Competing Interest Statement

The authors have declared no competing interest.

