## Supplementary information for "Donor transcription suppresses D-loops in *cis* and promotes genome stability"

### Author list and affiliations

### Supplementary figures and table legends

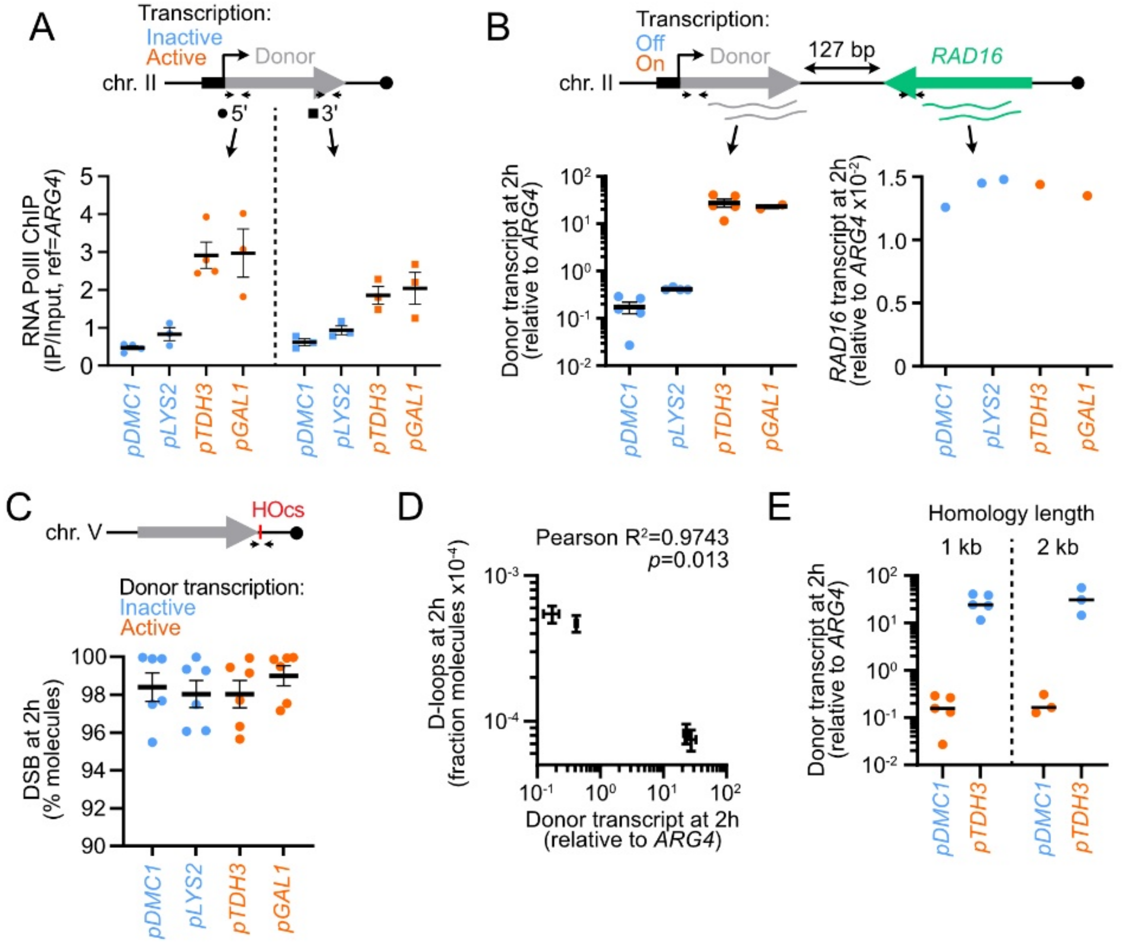


**Figure S1: Co-directional donor transcription suppresses nascent D-loops (related to Figure 1).**

A) ChIP-qPCR of the Rpb1 subunit of RNA PolII at the 5' and 3' end of the donor 2 hours post-DSB induction.

B) RT-qPCR of the RNA produced at the donor and at the downstream *RAD16* gene 2 hours post-DSB induction.

C) Quantification of the DSB frequency at *HOcs* 2 hours post-DSB induction.

D) Correlation between the D-loop levels and the amount of donor transcript. Data show mean ± SEM. From data in B) and **Fig. 1C**.

E) RT-qPCR of the RNA produced at the 1 kb- and 2 kb-long donors 2 hours post-DSB induction.

A-C, E) Data show individual biological replicates and mean ± SEM. * denotes statistical significance ($p<0.05$).

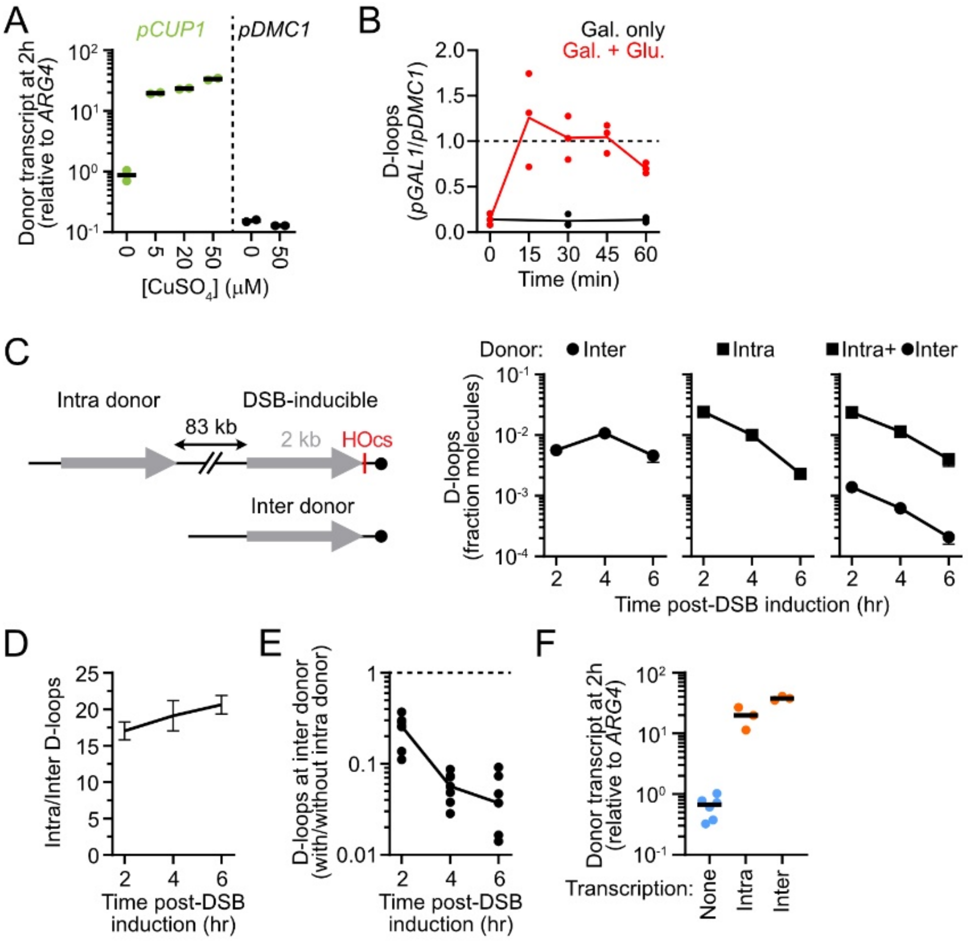


**Figure S2: Donor transcription suppresses D-loops in *cis* (related to Figure 2).**

A) RT-qPCR of the RNA produced at the donor with varying copper concentrations 2 hours post-DSB induction.

B) Ratio of D-loops formed at the donor site under the control of the *pGAL1* over the *pDMC1* promoter in contexts in which the *pGAL1* promoter is active (Gal. only) or at increasing time post-shutoff (Gal. + Glu.). From data in **Fig. 2B**.

C) Kinetics of D-loop levels at an intra-chromosomal and/or an inter-chromosomal donor (APY266, APY826, and APY809).

D) Ratio of D-loops formed at the intra donor over the inter donor when both are present (APY809). From data in C).

E) Fold decrease of D-loop at the inter donor when the intra donor is present. From data in C).

F) RT-qPCR of the donor RNA produced in strains bearing an intra and an inter donor either non-transcribed (APY809), with the intra donor transcribed (APY1587), or with the inter donor transcribed (APY1709).

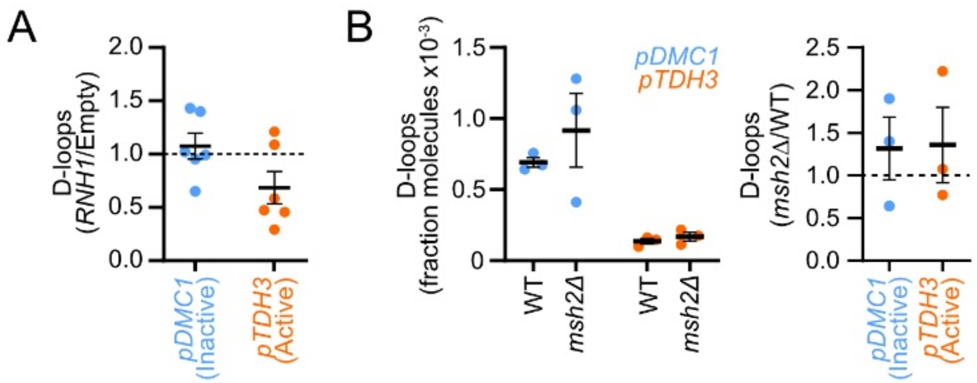


**Figure S3: Genetic determinants of transcription-mediated D-loop suppression (related to Figure 3).**

A) D-loops level fold change at transcriptionally inactive and active donors upon *RNaseH1* over-expression. From data in **Fig. 3A**.

B) Left: absolute D-loop levels at transcriptionally inactive and active donors in WT (APY502 and APY725), *msh2Δ* (APYand) strains. Right: relative mutant values compared to a WT strain assayed in parallel.

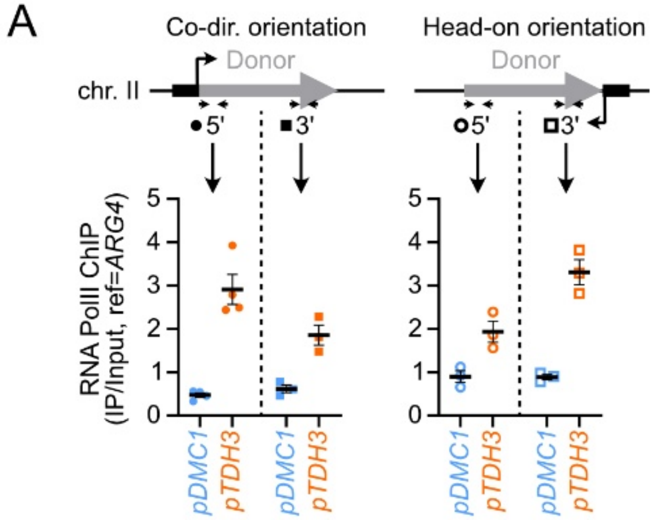


**Figure S4: Effect of transcription directionality (related to Figure 4).**

A) ChIP-qPCR of the Rpb1 subunit of RNA PolII at the 5’ and 3’ end of the donor 2 hours post-DSB induction with promoters in the co-directional or head-on orientation.
